# Downtown Diet: a global meta-analysis of urbanization on consumption patterns of vertebrate predators

**DOI:** 10.1101/2020.12.19.423628

**Authors:** Siria Gámez, Abigail Potts, Kirby L. Mills, Aurelia A. Allen, Allyson Holman, Peggy Randon, Olivia Linson, Nyeema C. Harris

**Author notes:** these authors contributed equally (Siria Gámez, Abigail Potts). Corresponding author: Siria Gámez.

## Abstract

Predation is a fundamental ecological process that shapes communities and drives long-term evolutionary dynamics. As the world rapidly urbanizes, it is critical to understand how the built environment and other human perturbations alter predation across taxa. We conducted a meta-analysis to quantify the effects of urban environments on three components of trophic ecology in predators: dietary species richness (DSR), dietary evenness (DEV), and stable isotopic ratios (δ^13^C and δ^15^N IR). We then evaluated whether intensity of anthropogenic pressure, using the human footprint index (HFI), explained variation in the effect sizes of dietary attributes using a meta-regression. We calculated Hedges’ g effect sizes from 44 studies including 11,986 samples across 40 predatory species in 39 cities globally. The direction and magnitude of effect sizes varied between predator taxonomic groups with reptile diets exhibiting the most sensitivity to urbanization. Effect sizes revealed that predators in cities had comparable DSR, DEV, and nitrogen ratios, though carbon consumption was significantly higher. We found that HFI did not explain variation in effect sizes, a result consistent between the 1993 and 2009 editions of this metric. Our study provides the first assessment of how urbanization has perturbed predator-prey interactions for multiple taxa at a global scale, revealing that the functional role of predators is conserved in cities and urbanization does not inherently relax predation.

## Introduction

Predation is a process which underpins ecological and evolutionary dynamics at various scales, from the individual to the ecosystem. Predation can increase regional species richness and diversity by mediating competition in prey species (Holt, 1977; Shurin, & Allen, 2001). Moreover, predators alter ecosystem-level processes such as nutrient cycling by provisioning carcasses and enriching soil or water columns (Schmitz et al., 2010). Apart from consumptive effects, predation can structure communities indirectly through trophic cascades (Terborgh, 2010). The fear of predation itself can engender non-consumptive effects that alter space use and aggregation of prey and subsequently drives vegetation patterns (Suraci et al., 2016; Gaynor et al., 2019). While the theoretical and empirical literature is rich with studies quantifying the effects of predation in natural systems, our understanding of how urban environments affect predation remains limited, even contradictory. For example, predation rates on human landscapes can be amplified by increased prey densities or relaxed due to an abundance of easily accessible anthropogenic subsidies, creating an urban predation paradox (Rodewald et al., 2011; Fischer et al., 2012).

Cities are an emerging socio-ecological ecosystem inducing novel interactions, behavioral shifts, and evolutionary trajectories (Lowry et al., 2013; Johnson, & Munshi-South, 2017; Lepczyk et al., 2017). By 2030 more than 60% of the world’s human population is projected to live in an urban area (United Nations, 2018). The effects on the landscape from such rapid urbanization are profound; globally, urban land cover is projected to increase by 1.2 million km^2^ by 2030, decimating available habitat for wildlife and reducing agricultural land by 550,000 km^2^ (Seto et al., 2012; Bren D’amour et al., 2017). Urbanization can decrease prey species richness and genetic diversity, alter community composition, and prey body size; thus, altering resource availability and diet selection for secondary and tertiary consumers (El-Sabaawi, 2018; Chejanovski, & Kolbe, 2019; Schmidt et al., 2020). Regional species pools are further filtered by urban form and history, novel urban species interactions, and disparate distributions of natural resources in the urban landscape due to systemic racism (Aronson et al., 2016; Schell et al., 2020). Additionally, human food subsidies increase trophic niche overlap in terrestrial carnivores, potentially resulting in greater interspecific competition (Manlick, & Pauli, 2020). Cities also modify wildlife behavior, which influences vulnerability to predation or access to food, by disrupting diel patterns and vigilance behaviors (Gaynor et al., 2018; Gallo et al., 2019).

Urbanization is a complex anthropogenic process that can alter predator-prey interactions through a multitude of potential mechanisms associated with changes to food availability, habitat connectivity, vegetation density, and microclimate (Alberti et al., 2020; Johnson et al., 2020). Anthropogenic infrastructures bisect habitat, increase the cost and mortality risk of movement, and create novel temperature gradients, driving changes to population and community-level processes including predation (Golden, 2004; Fraser et al., 2019; Zambrano et al., 2019). Habitat fragmentation can reduce prey abundance, affecting predator diet selection and evenness (Layman et al., 2007). In particular, roads cause a significant proportion of wildlife mortality, upwards of 49% of all adult and juvenile mortality for some species, and underlying the trend of death rates exceeding birth and recruitment rates in urban areas (Bateman, & Fleming, 2012; Tucker et al., 2018). In addition to fragmentation, extensive homogenization of urban vegetation structure can reduce overall cover and affect prey behavior and space use (Denno, & Kaplan, 2007).

The effects of such varied mechanisms induced from urbanization manifest differently among taxonomic groups with wildlife responses being scale-dependent (Fidino et al., 2020). For example, cities exhibit extreme temperature gradients, which could result in varied consumptive patterns, as low temperatures increased attack rate in *Daphnia*, while higher temperatures increased the prey consumption rate in reptiles and birds (Wasserman et al., 2016; Scott et al., 2017). Herpetofauna are more susceptible to higher disease prevalence and toxicity in urbanized ecosystems compared to mammal fauna (Murray et al., 2019). In some cases, urbanization can even hamper the spread of disease as a result of reduced host densities in cities compared to rural areas (Fountain-Jones et al., 2017; Gras et al., 2018). While patterns of species richness and population density vary significantly across taxa, urban birds and arthropods tend towards reduced diversity and increased abundance (Faeth et al., 2011; Mcdonald et al., 2020).

Despite the recent surge of urban ecology studies employing comparative urban vs. non-urban frameworks, broad scale predation patterns across taxa remain largely unknown. Studies often focus on a single species or city, limiting inference at a broad scale. Additionally, a lack of a standardized definition of “urban” has made cross-city comparisons challenging, coupled with varied experimental designs, sample sizes, and bias towards readily observable study organisms. Thus, we lack a systemic understanding of how the urban environment affects predator diet, how it varies across taxa, and how these effects scale with the intensity of human impact on the landscape. Further, assumptions regarding the relaxation of predation in urban spaces remain untested across multiple taxonomic groups. Here, we conduct the first global meta-analysis of how urban environments affect three aspects of predator trophic ecology: dietary species richness (DSR), dietary evenness (DEV), and trophic niche using δ^13^C and δ^15^N isotopic ratios (IR) (Figure 1). Specifically, we address the following questions:

1. How does urbanization affect predator diet composition? We expect a decrease in DSR, given documented reductions in species richness due to anthropogenic perturbations in urban environments (Ortega-Álvarez, & Macgregor-Fors, 2009; Ferenc et al., 2014). If species richness in cities declines, the relative abundance of some common urban prey species can increase significantly. Therefore, we expect DEV to decrease in urban areas (Mccabe et al., 2018). We expect the urban predator trophic niche to reflect lower δ^15^N and greater δ^13^C ratios relative to their rural counterparts due to exploitation of anthropogenic food sources rich in corn, wheat, and sugar (Murray et al., 2015; Manlick, & Pauli, 2020; Scholz et al., 2020).
2. How do the effect sizes of urbanization on dietary attributes differ across predator taxa? We expect trophic responses to urbanization to differ significantly among predator taxa, owing to implicit differences in diet plasticity, behavior, and natural history as well as biased representation of taxa in urban ecology literature (Bonnet et al., 2002; Samia et al., 2015). Such variation in sensitivity to perturbations in urban ecosystems may ultimately drive heterogeneity in the direction and significance of effect sizes among wildlife taxa (French et al., 2018; Rocha, & Fellowes, 2018).
3. How do urban effects on predation relate to the human footprint index (HFI)? We expect the effect sizes to positively correlate with the HFI, indicating that as the intensity of anthropogenic pressure increased, the magnitude of change to predation would increase. Predator diversity and density, and thus predation rates and prey selection, are correlated with the components used to calculate the HFI, such as human population density and land-use conversion, underpinning our expectation that the degree of urbanization would explain variation in predator diet effect sizes (Eötvös et al., 2018; Korányi et al., 2020).

**Figure 1:**
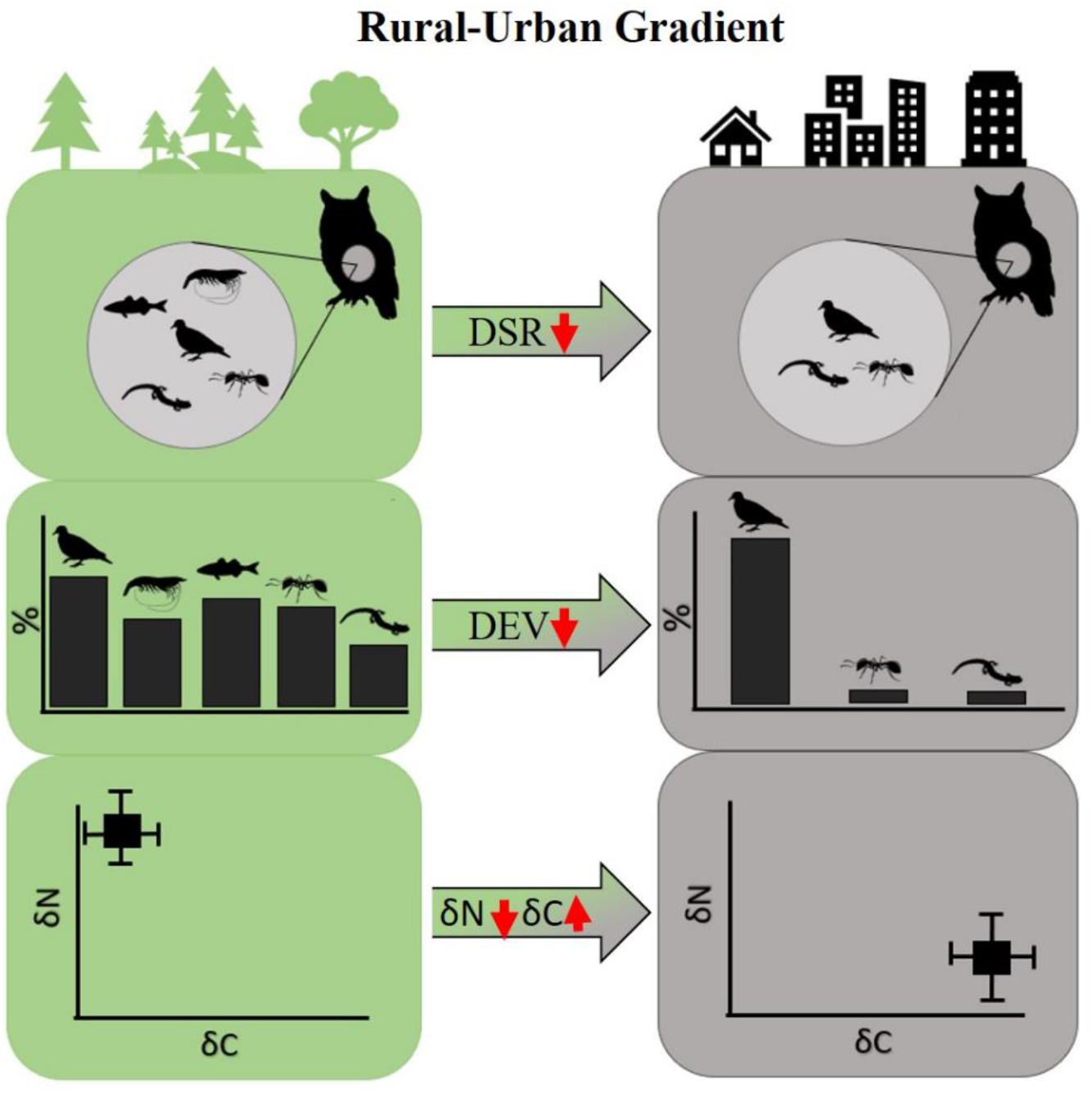
Conceptual diagram illustrating how cities can influence three components of predator trophic ecology, a) dietary species richness (DSR), b) dietary evenness (DEV), and c) δ^13^C and δ^15^N isotopic ratios. Green (left) column represents rural and wildland habitat, while grey (right) denotes urban habitat.

## Methods

### Literature review

To quantify the effects of urbanization on predation, we completed a comprehensive literature search of empirical studies that provided estimates on aspects of predation rate, prey availability vs selection, or prey diversity in the context of a predator of any taxa and compared these dietary metrics in an urban vs. rural or wildland framework (discrete or gradient). First, we conducted a broad topic search using the Web of Science database with the following terms: “predat* AND urban”, “prey AND urban”, “prey AND urban AND rural”. We then conducted a second, stricter search by filtering results by topic “Ecology”, “Zoology”, “Biodiversity Conservation”, “Environmental Sciences”, or “Ornithology”, using the keywords: “resource AND use AND urban”, “predat* AND urban AND wildland”, and “carniv* AND urban AND diet”. We did not limit the inclusion of studies based on time of publication if they satisfied our other criteria. Searches were conducted iteratively (Figure S1); we recorded the number of results for each search iteration following the Preferred Reporting Items for Systematic Reviews & Meta-Analyses (PRISMA) guidelines (Moher et al. 2015).

Without a unified definition of “urban” or use of standardized response variables in the urban ecology literature, we defined the selection criteria for inclusion in the analysis. Prospective studies needed to report summary data of predator species diet or predation metrics including mean, variance, and/or standard deviation for both urban and rural categories to calculate effect sizes. How studies defined urban and rural varied considerably. For example, some studies used a categorical approach compared to a continuous gradient of urbanization based on an index such as % impervious surface, human population, or distance to the urban core. In studies that sampled urban vs. rural in a discrete fashion, we simply extracted values for each category. For studies that used a gradient approach, we extracted predation and prey composition values from the two extremes.

### Predation consumption metrics

We explored three metrics of trophic ecology in our analysis to capitalize on the multiple methods presented in the literature. We extracted DSR data by counting each prey taxon observed in the predator’s stomach contents, pellets, or through direct predation observations. Species richness in a predator’s diet reflects dietary breadth. We also quantified DEV by calculating the standardized niche breadth (Eq. 1) for urban and rural samples in each study (Newbury, & Hodges, 2018). Evenness contrasts with species richness in that the relative frequency and representation of each prey type is considered (Levins, 1968; Formoso et al., 2012). We recorded sample sizes and sample standard deviations where possible. However, ~90% of included studies had no associated measure of within-study variance because the studies did not all use DEV as a response variable for individual samples. To address this limitation, we used a maximum likelihood estimation (MLE) and a random effects model to estimate heterogeneity for each study’s observed mean difference, enabling us to fill in missing variance values (Sangnawakij et al., 2017).

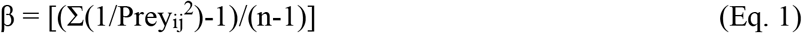

Carbon and nitrogen isotopic ratios present in scat, hair, and vibrissae allow researchers to assess individual heterogeneity in the trophic niche space (Ben-David, & Flaherty, 2012; Colborn et al., 2020). Higher δ^13^C IR reflect consumption of plants with C4 photosynthetic pathways such as corn and sugarcane, which are common in anthropogenic food sources, in contrast to lower values associated with C3 plants found in rural or wildland habitat (Galetti et al., 2016). High δ^15^N IR indicate consumption of protein-rich animal prey, denoting trophic status and degree of carnivory (West et al., 2006; Marshall et al., 2019). Because studies generally depict present δ^13^C and δ^15^N isotopic ratio values graphically in a 2-dimensional isoscape, we extracted data of urban and rural samples from figures using the program WebPlotDigitizer (Rohatgi, 2015). We used all three components of trophic ecology, DSR, DEV, and IR, as a distinct response variable in our meta-analysis to isolate individual responses in predators to changes in urbanization.

### Statistical analysis

We calculated the effect size of each study using the *Hedges’ g* metric that includes a correction term for small sample sizes to determine consequences of urbanization on predation metrics. (Cooper, 1994). Positive effect sizes indicate an increase in the response variable, such as a greater DEV in urban environments compared to the rural/wildland control. Between-group heterogeneity τ^2^ was estimated using the Sidik-Jonkman method and assumed to be equal for all predator taxonomic groups; larger τ^2^ values indicate greater variance of observed effect sizes between taxa (Sidik, & Jonkman, 2007). Effect sizes were derived for each individual study, for grouped predator taxa, and across all studies. Latitude and longitude coordinates were collected from each study’s urban and rural/wildland sampling sites. For studies where exact points were not reported, we extracted coordinates based on the centroid of the study areas for discrete sites. For sites where urban-rural gradient transects were employed, we assigned coordinates at the extreme points of the studies transects. (i.e. urban core versus most peripheral sampling point). We then extracted HFI values for urban and rural sites within each corresponding study using ArcMap (ArcGIS Desktop v 10.7). The HFI, whose values range from 0 to 50, is an effort to quantify anthropogenic pressures on the landscape by incorporating built environments, human population density, electric infrastructure, crop lands, pasture, roads, railways, and navigable waterways into a single metric at a global scale (Venter et al., 2016).

We used mixed effect meta-regression models to determine how predator taxa and ΔHFI explain the variability of observed effect sizes (Fleiss, 1993). We calculated ΔHFI as HFI_URBAN_-HFI_RURAL_, where positive values indicate greater anthropogenic pressures as expected in the urban site. We derived two versions of ΔHFI using the metric as it was calculated in 1993 and later in 2009 (Venter et al., 2016). Given the range in publication dates among papers included in our study (1986-2020), we repeated the meta-regression for both versions of ΔHFI to determine whether results were robust to when the metric was calculated. We derived a regression coefficients (β) and 95% confidence intervals for each taxonomic group to determine whether ΔHFI explained variation in effect sizes. Significance was determined based on whether the β and its confidence interval did not overlap zero. To test for publication bias in the studies included in the analysis, we built a funnel plot to visualize the overall spread of effect sizes and their corresponding error estimates (Figure S2). Finally, we performed Kendall’s Rank test to quantitatively assess the correlation between effect size and error estimates. The meta-analyses were carried out in program R (v. 3.6.3) using the ‘*metafor*’ and ‘*meta*’ packages in Program R (R Core Team, 2017).

## RESULTS

Our initial search yielded 358 studies related to predation in urban versus rural or wildland systems. Based on our selection criteria, 32 of these studies were included in our final analysis. We performed a subsequent targeted search to improve representation for amphibians and reptiles; this resulted a total of 62 potential studies of which 9 satisfied the criteria for inclusion in our analysis. Our final analysis represented a total of 44 studies with 57 effect sizes calculated that spanned across 39 cities in 6 continents (Figure 2). One study reported diet for two species and another study reported both diet content and stable isotope ratios. Our synthesis yielded 11,986 samples of predator stomach contents, predation events, hair, and pellets across 40 species from 5 predatory taxa. Mammals (n=19) and birds (n=15) had the greatest representation of studies included in our analysis, with 43.2% and 34.1%, respectively. Fish (n=4) and reptiles (n=4) each comprised 9% of represented studies, while amphibians (n=2) made up 4.5%. Terrestrial and aquatic systems comprised 88.6% and 11.4% of included studies, respectively.

**Figure 2:**
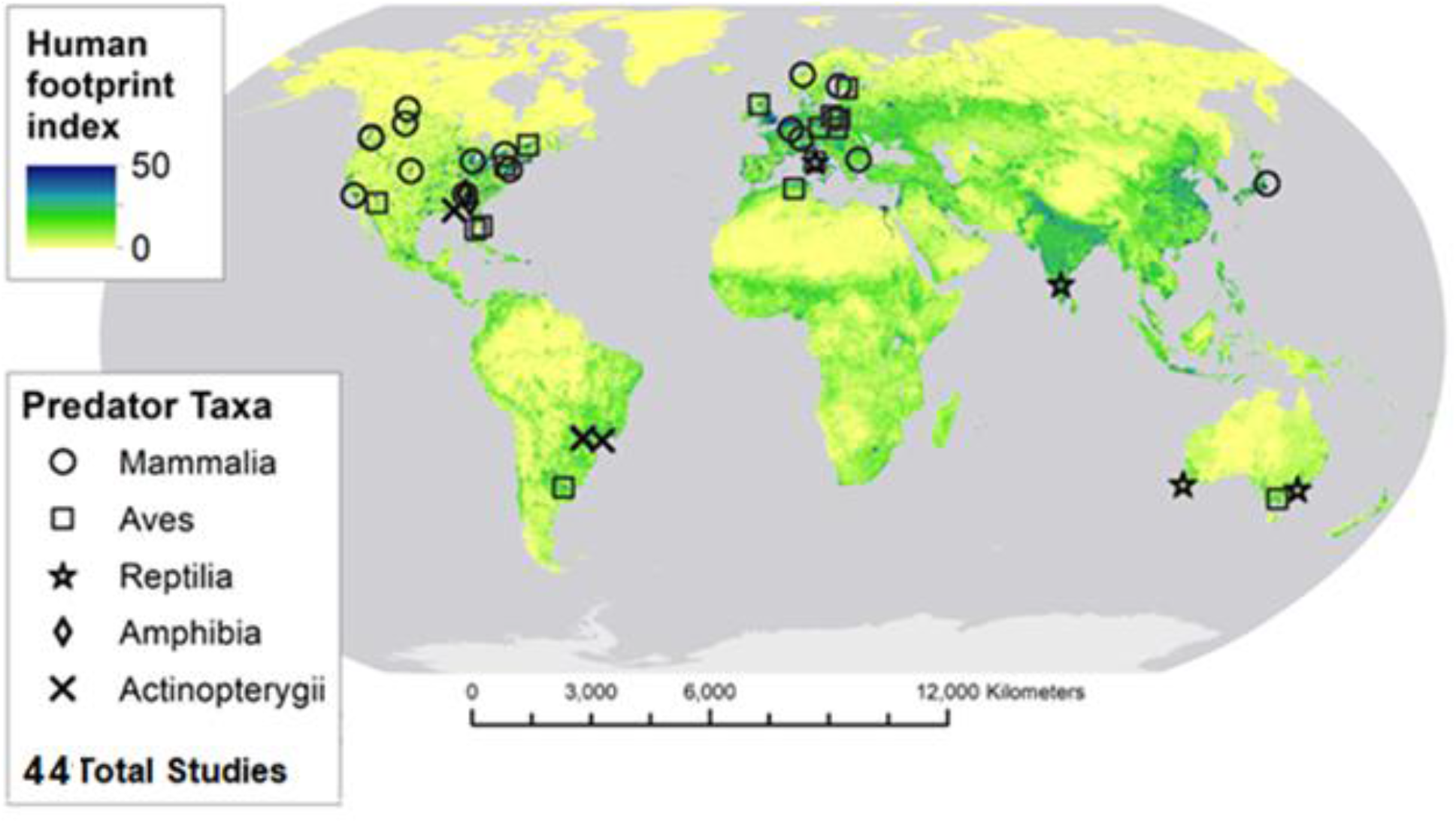
Global distribution of studies included in the analysis. Symbols represent predator taxa (class), and color gradient illustrates human footprint index (HFI) from 2009.

### Predation consumption metrics

We investigated consequences of urbanization on three different predation consumption metrics that represent a species’ trophic ecology. Contrary to our hypothesis, overall DSR was not significantly lower in urban environments when studies were aggregated (Hedges’ g: −1.74, τ^2^ = 149.08, 95% CI: −6.08, 2.60). The direction of the effect varied between predator taxa, though the effect size and respective 95% confidence interval overlapped zero for all taxa (Figure 3a). Fish, bird, and reptile DSR decreased in response to urbanization, indicating a more specialized foraging strategy. In contrast, mammals and amphibians consumed a greater number of prey species and adopted a more generalist foraging strategy in urban environments.

**Figure 3:**
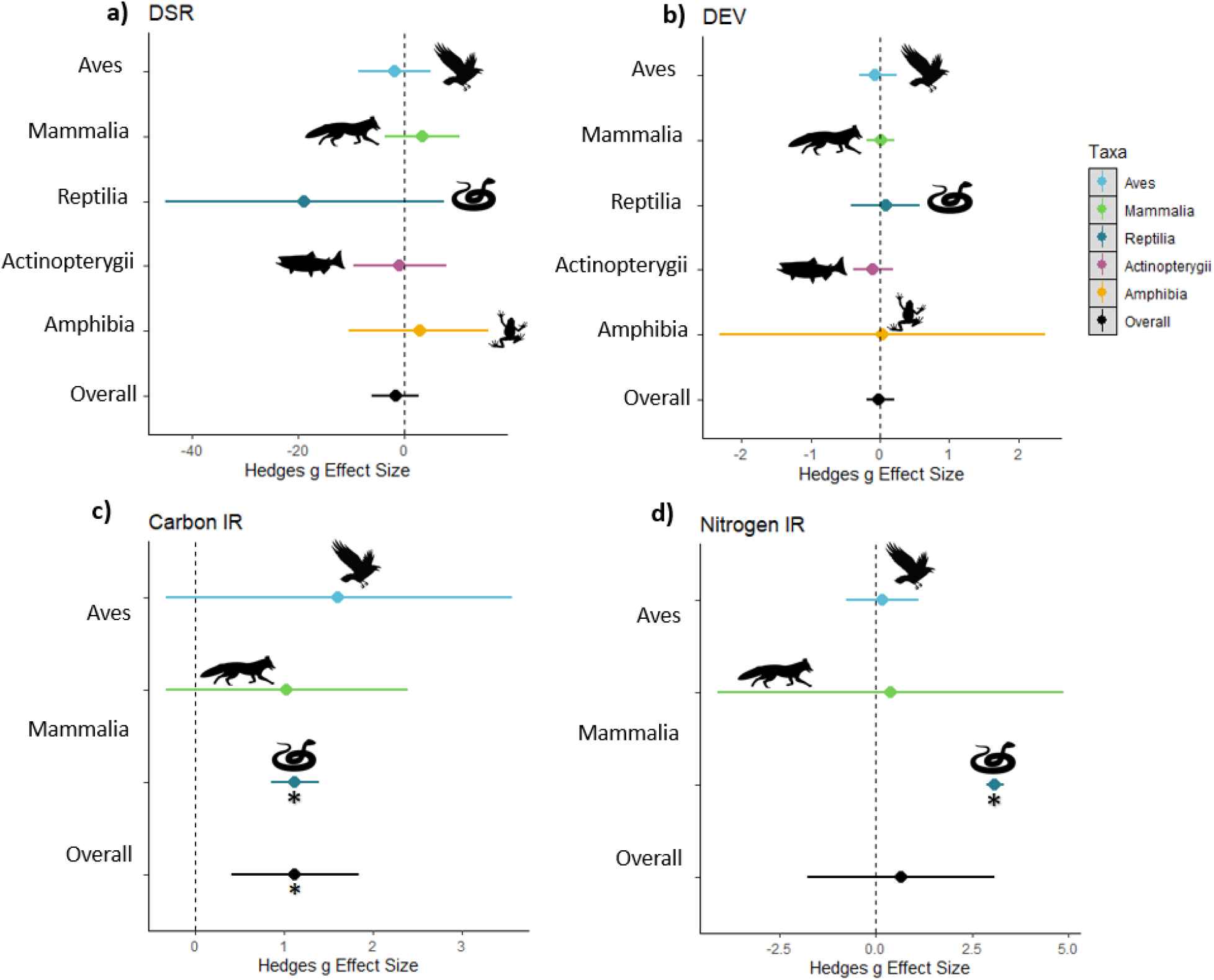
Effect sizes for components of trophic ecology grouped by taxa: a) dietary species richness – DSR; b) dietary evenness - DEV, c) δ^13^C and d) δ^15^N. Hedge’s g used to calculate effect sizes and asterisk indicates significant effect of urbanization on diet metric based on whether 95% confidence interval overlaps zero. ΔHFI 2009.

Contrary to expectations, overall DEV was also not significantly lower in cities compared to rural or wildland areas (Hedge’s g: −0.02, τ^2^ = 0.02, 95% CI: −0.08, 0.04). Reptiles, mammals, and amphibians showed greater DEV (i.e., comparable representation of individual prey items) in cities (Figure 3b) while, bird and fish diets were skewed in urban environments, meaning a relatively small number of prey types dominated consumptive patterns, though effect sizes and 95% confidence intervals overlapped zero for all taxonomic subgroups.

We found evidence that diets of urban predators were more carbon-enriched, as evident by significantly higher δ^13^C isotopic ratios compared to rural predators (Figure 3c-d; Hedge’s g: 1.12 τ^2^ = 0.65, 95% CI: 0.41, 1.84). In contrast, urban δ^15^N ratios were not significantly different than rural or wildland predators (Hedge’s g: 0.67 τ^2^ = 8.42, 95% CI: −1.76, 3.10). Overall, urbanization had a stronger influence on δ^15^C ratios, signaling that predators are have adopted a strategy of consuming vast quantities of anthropogenic food sources in cities, rich in sugar, corn, and wheat.

### Human Footprint Index

Values of HFI varied greatly, even among urban sites, highlighting the breadth of intensity of anthropogenic pressures across “urban” areas. The average HFI for urban sites was 28.7 (range = 17.3 - 49.4) while only 17.8 (range = 1.1-48.4) for rural sites in 1993 dataset and In comparison, anthropogenic influences estimates were higher in 2009 with the average HFI for urban sites increasing 1.5% (mean = 29.1; range= 18-49.4), while the average HFI for rural sites increasing 13% (mean = 20.1; range=1.3-48.4). When total effect sizes for all predator taxa were pooled, ΔHFI_2009_ had a non-significant relationship to the effect of urbanization on all consumption metrics (DSR: β = 0.029, 95% CI = −0.02, 0.08; DEV: β = −0.002, 95% CI = −0.19, 0.15; δ^13^C: β = −0.003, 95% CI = −0.31, 0.29; δ^15^N: β = −0.002, 95% CI = −2.1, 2.18). Considering how DEV responded by taxa, we found that bird DEV had a significant, negative response to ΔHFI_2009_ (β = −0.004, 95% CI = −0.008, −0.00008). Conversely, fish DEV had a significant, positive response to ΔHFI_2009_ (β = 0.007, 95% CI = 0.003, 0.01), though overall DEV effect size response to ΔHFI_2009_ was non-significant. For DSR as well as carbon and nitrogen IR, ΔHFI_2009_ did not explain variation in effect sizes. Results using ΔHFI_1993_ mirrored those of the 2009 metric with non-significant relationships in the effect of urbanization across consumption metrics. Therefore, we found little evidence that the degree of anthropogenic change on the landscape significantly influenced the magnitude of effect sizes of urbanization on predation (Figure 4).

**Figure 4:**
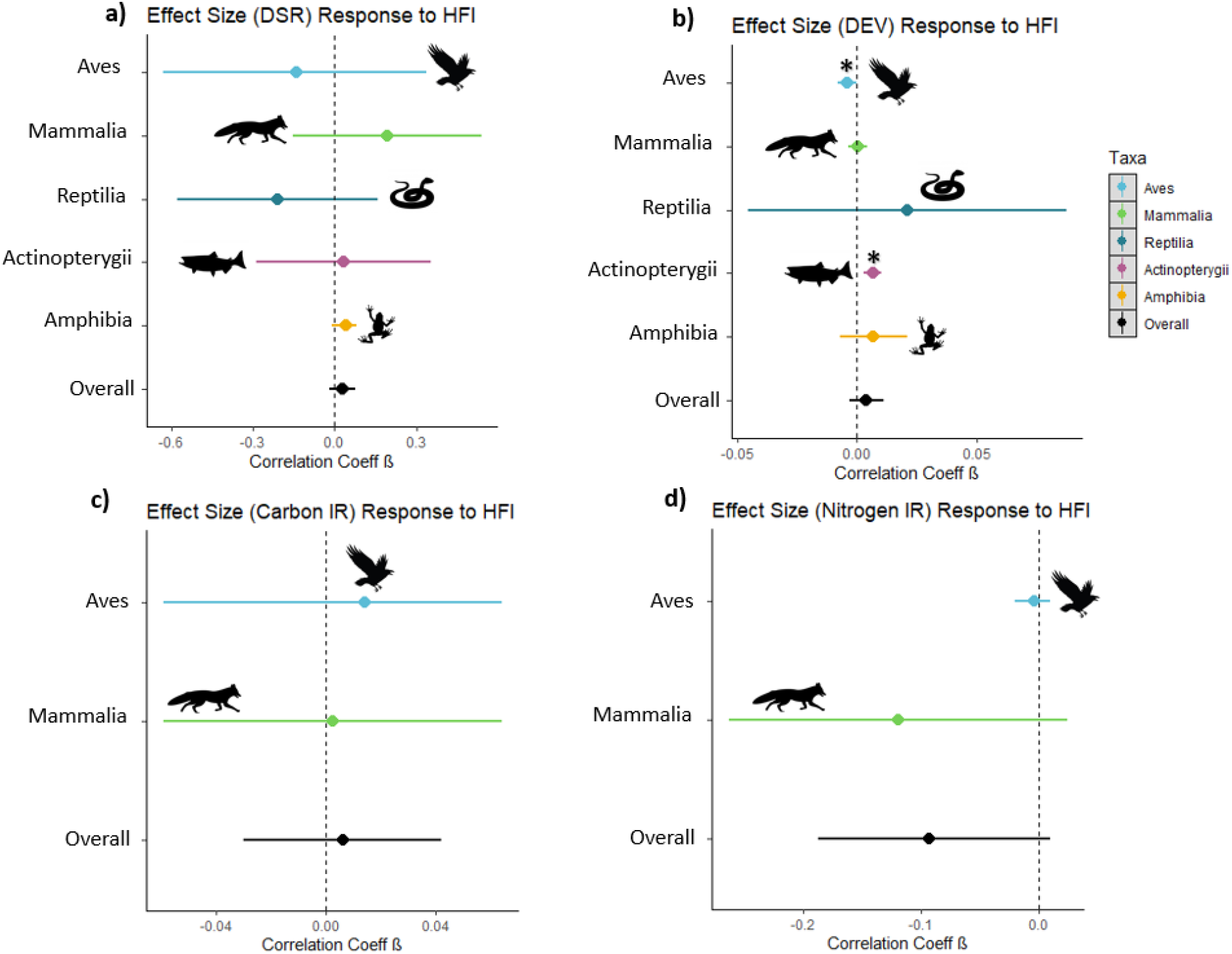
Meta-regressions of effect size response to change in human footprint index from rural-urban sites from 2009 (ΔHFI), a) DSR, b) DEV, c) δ^13^C, and d) δ^15^N

## DISCUSSION

As urbanization alters landscapes worldwide, it is critically important to understand and anticipate the ecological consequences to wildlife living within the built environment (Alberti et al., 2017; Aronson et al., 2017). Our global meta-analysis revealed that predator trophic ecology changed significantly in consumption of carbon, but was conserved for other. Alarmingly, taxonomic groups such as amphibians, which are poorly represented in the urban ecology literature and our meta-analysis, are also experiencing disproportionately high extinction rates (Alroy, 2015). Therefore, global regions with high species-richness coupled with rapid urbanization are particularly at-risk of extensive changes to its faunal community composition, underscoring the importance of comparative urban-rural studies on poorly studied taxa.

Broad-scale shifts to DSR and DEV in predator diet carry profound ecological implications for predator-prey relationships, population regulation, and disease transmission. The decline in prey species richness could increase dietary overlap for interspecific competitors, resulting in competitive exclusion (Du Preez et al., 2017; Manlick, & Pauli, 2020). Fewer prey species in cities promotes higher population densities for those species that can exploit the urban environment (Steinberg et al., 1997; Uno et al., 2010). Such surges in prey population densities can in turn drive predator population increases, potentially fueling human-wildlife conflict (Yirga et al., 2017; Fleming, & Bateman, 2018; Mccabe et al., 2018).

Diets of urban wildlife are not necessarily “protein poor”, as evidenced by comparable levels of nitrogen detected in isotopic signatures. However, consumption of carbon-based food items increased in cities, likely as a result of corn, wheat, and sugar-rich anthropogenic refuse (Bateman, & Fleming, 2012) characteristic of urban habitats. Our results highlight that the ecological role of predators as top-down regulators is conserved, not relaxed in cities. Abundant prey species associated with urban environments such as brown rats (*Rattus norvegicus*), rock pigeons (*Columba livia*), and house mice (*Mus musculus*) could potentially be driving predation rates and protein (i.e. nitrogen) consumption, at least for mammalian and avian predators (Corsini et al.; Newsome et al., 2010; Mccabe et al., 2018; Scholz et al., 2020). Our findings are contrary to an emerging hypothesis of relaxed predation phenomenon in urban areas, described by a meta-analysis of 25 studies that found predation rates on bird nests were reduced in urban areas (Eötvös et al., 2018).

Effect sizes did not correlate with the change in human footprint index, meaning the intensity of anthropogenic alterations to the landscape did not significantly amplify observed changes to predator’s diet. These results were robust to the year when HFI was calculated, signifying that perturbations to predator trophic ecology had likely already occurred by the time HFI was first measured in 1993 and did not change drastically after the 2009 HFI census. A possible explanation for this result is that while urban ecosystems do induce changes to predator diet, these effects occur early in the urbanization process and do not continue to amplify as cities become denser and more developed. Many cities reported low ΔHFI values, meaning the difference in their urban core and rural site was low or close to zero. Importantly, HFI values for some “rural” sites were as high as those designated “urban” in other studies, underscoring a fundamental challenge in urban ecology for comparative works. Moreover, this overlap in HFI between rural and urban sites demonstrates that “rural” does not equate to “natural” or habitats devoid of anthropogenic perturbations. Rural areas encompass agricultural production such as crops and livestock, each known to alter vegetation and animal communities (Gordon, 2018; Lark et al., 2020). Historically, “urban-rural” comparative studies have framed these categorizations as dichotomous spaces and carried an implicit assumption about the relatively intact character of “rural” areas, which is not necessarily consistent at large spatial and temporal scales (Moll et al., 2019). Further, the HFI metric is a composite indicator of human pressures and includes both the built environment and agricultural activities, contributing to low ΔHFI values in comparisons between rural and urban sites.

While our results provide key insights on predation in urban environments, we recognize limitations that require future work. We found bias in taxonomic representation with more studies on predation across urban-rural gradients for birds and mammals. Taxonomic bias is a well-known trend in the conservation ecology field, where charismatic vertebrate species have been historically overrepresented in published studies (Griffiths, & Dos Santos, 2012). We also found bias in the distribution of sample sizes across taxa. Molecular techniques in predation studies commonly use pellets or scat, requiring additional analysis to identify individuals in the population. Fewer than 10% of included studies identified individual host identity, potentially skewing our interpretation of the effect of urbanization on predation and limiting our inference at broad ecological scales (Troudet et al., 2017). The geographic representation of biomes in published studies was largely skewed to terrestrial ecosystems, highlighting the urgent need to study aquatic systems in proximity to urban areas. Finally, given inconsistencies in studies reporting sample variances, we highlight the need for such estimates to facilitate future cross-taxa and cross-site comparisons of urban ecology.

Wildlife must increasingly adapt to city living and our synthesis underscores how the built environment modifies a fundamental ecological process. In a rapidly urbanizing world, perturbations to predator-prey relationships can drastically change human-wildlife interactions, ecosystem processes, and species extinctions, it is therefore critically important to understand these changes to inform future efforts to mitigate them. We recommend future studies aim to obtain long-term diet data at multiple sites for predatory species including at the individual level, as quantifying dietary changes over time with growing infrastructures can provide guidance on coexistence strategies for humans and wildlife (Newsome et al., 2015; Teyssier et al., 2020). The inclusion of stable isotopes in future urban-rural diet analyses would fill important information gaps to couple trophic and urban ecology, as these data capture a crucial dimension of niche space at a broader trophic level. Recent work has demonstrated variation in dietary niche and trophic position for urban versus rural coyotes using stable isotopes (Colborn et al., 2020). To conclude, we provide the first quantification of predator diets in a comparative urban versus rural or wildland framework for multiple predator taxa at a global multi-city scale and revealed that urbanization enriches predator diets with carbon but does not inherently relax predation.

## Acknowledgements

Our sincere thanks to members of the Applied Wildlife Ecology (AWE) Lab at the University of Michigan, specifically, R. Malhotra, S. Lima, and N. Arringdale who assisted with data collection and provided feedback on the conceptual framework and figures in this work. We also thank Drs. A. Ostling and N. Carter for their constructive comments which improved the manuscript.

